# Hierarchical chromatin organization detected by TADpole

**DOI:** 10.1101/698720

**Authors:** Paula Soler-Vila, Pol Cuscó Pons, Irene Farabella, Marco Di Stefano, Marc A. Marti-Renom

**Affiliations:** CNAG-CRG, Centre for Genomic Regulation (CRG), Barcelona Institute of Science and Technology (BIST), Barcelona, Spain; Gastrointestinal and Endocrine Tumors Group, Vall d’Hebron Institute of Oncology (VHIO), Barcelona, Spain; Centre for Genomic Regulation (CRG), Barcelona Institute of Science and Technology (BIST), Barcelona, Spain; Universitat Pompeu Fabra (UPF), Barcelona, Spain; ICREA, Barcelona, Spain

## Abstract

The rapid development of chromosome conformation capture (3C-based) techniques as well as super-resolution imaging together with bioinformatics analyses has been fundamental for unveiling that chromosomes are organized into the so-called topologically associating domains or TADs. While these TADs appear as nested patterns in the 3C-based interaction matrices, the vast majority of available computational methods are based on the hypothesis that TADs are individual and unrelated chromatin structures. Here we introduce TADpole, a computational tool designed to identify and analyze the entire hierarchy of TADs in intra-chromosomal interaction matrices. TADpole combines principal component analysis and constrained hierarchical clustering to provide an unsupervised set of significant partitions in a genomic region of interest. TADpole identification of domains is robust to the data resolution, normalization strategy, and sequencing depth. TADpole domain borders are enriched in CTCF and cohesin binding proteins, while the domains are enriched in either H3K36me3 or H3k27me3 histone marks. We show TADpole usefulness by applying it to capture Hi-C experiments in wild-type and mutant mouse strains to pinpoint statistically significant differences in their topological structure.

## INTRODUCTION

The organization of the genome in the cell nucleus has been shown to play a prominent role in the function of the cell. Increasing evidence indicates that genome architecture regulates gene transcription (1,2) with implications on cell-fate decisions (3–5), development (6), and disease occurrences such as developmental abnormalities (7,8) and neoplastic transformations (9–11).

The genome organization is characterized by complex and hierarchical layers (1). For example, fluorescence *in-situ* hybridization revealed that chromosomes are positioned in preferential areas of the nucleus called chromosome territories (12). This large-scale feature has been confirmed by high-throughput Chromosome Conformation Capture (Hi-C) experiments (13), that provided a genome-wide picture in which inter-chromosomal interactions are depleted relative to intra-chromosomal. Analysis of Hi-C data also revealed the segregation of the genome in multi-megabase compartments characterized by different GC-content, gene density, and chromatin marks (13–15). Microscopy approaches, in spite of considerable variability, have corroborated the spatial segregation of such compartments at the single cell level (16). At the sub-megabase level, Hi-C experiments also revealed the presence, validated by microscopy approaches (17–19), of self-interacting regions termed Topologically Associated Domains (TADs) (20,21). TADs are composed by dense chromatin interactions, which promote 3D spatial proximity between genomic *loci* that are distal in the linear genome sequence. Since many of these interacting *loci* are cis-regulatory elements, TADs are usually considered as the structural functional units of the genome that define the regulatory landscape (22,23), and are conserved across cell types and species (20,24). Moreover, TADs boundaries are often demarcated by housekeeping genes, transcriptional start sites and specific chromatin insulators proteins, such as CTCF factor and cohesin complex (20,25). TADs appear to be further organized in a hierarchical fashion. For example, in mammalian cells, concepts such as “metaTADs” (26) or “sub-TADs” (27) have been introduced. The former is used to define a superior hierarchy of domains-within-domains that are modulated during cell differentiation (26) while the latter to emphasize how and where the cis-regulatory elements establish physical interactions that contribute to gene regulation (27).

Several computational methods to identify and characterize TADs from 3C-based interaction data have been reported (28,29). Based on different *a priori* assumptions on the TADs subdivision, these methods can be mainly classified as disjointed or overlapping. The former consider TADs as individual and unrelated structures with no possible mutual intersections (*e.g*. directionality index (DI) (20), insulation score (IS) (30), ClusterTAD (31), ICFinder (32)). The latter assume that TADs are overlapping and related structures with a shared content (e.g. Arrowhead (13,15), armatus (33), TADtree (34), 3DNetMod (35)). However, only few algorithms (CaTCH (36), GMAP (37), matryoshka (38), and PSYCHIC (39)) can identify nested domains where each domain contain other sub-domain profiling a hierarchical chromatin architecture.

Here, we present TADpole, a bioinformatics tool to disentangle the full structural chromatin hierarchy that automatically determines an optimal division level. Notably, TADpole is robust both at technical and biological benchmarks based on a published study (29) and does not rely on mandatory parameters. We prove the effectiveness of TADpole to investigate the chromatin hierarchy in capture Hi-C data (cHi-C) (40) where the chromosome topology is altered with local genomic inversions that drive gene misexpression associated to congenital malformations in mouse (41).

## MATERIAL AND METHODS

### The TADpole pipeline

TADpole consists in three main steps (**Figure 1A**): (*i*) pre-processing of the input Hi-C dataset, (*ii*) constrained hierarchical clustering optimization, and (*iii*) genome segmentation. TADpole has been implemented as an R package available at https://github.com/3DGenomes/TADpole.

**Figure 1.**
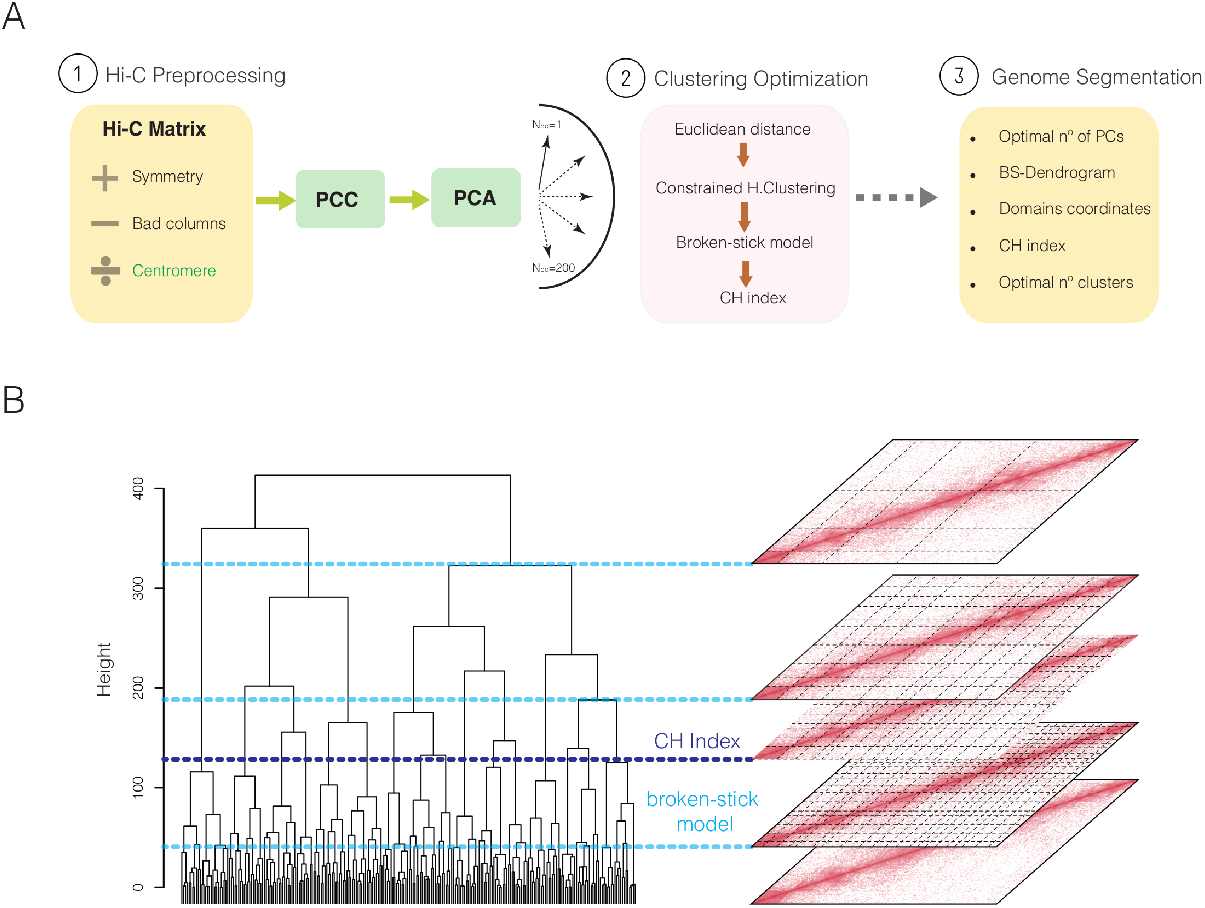
General overview of TADpole tool. **(A)** Schematic view of TADpole algorithm. (1) TADpole input is an all-vs-all interaction matrix. The matrix is checked for symmetry, and low-quality columns (bad columns) are removed. Large matrices of entire chromosomes are optionally split at the centromere to create two smaller sub-matrices corresponding to the chromosomal arms. Next, denoising and dimensionality reduction steps take place by computing the corresponding Pearson’s correlation coefficient (PCC) matrix, and its principal component analysis (PCA). (2) Per each number of first PCs retained (from 2 to 200), the PC matrix is transformed into the corresponding Euclidean distance, that serves as input to perform the constrained hierarchical clustering. The range of possible clustering levels in the hierarchy is given by an upper bound according to a broken-stick model, and then the Calinski-Harabasz (CH) index is used to select the optimal partition. (3) As output, TADpole returns the number of first PCs retained to obtain the optimal set of TADs, the dendrogram with the significant hierarchical levels, the coordinates of the chromatin domains for each partition with the associated CH index, and the optimal number of clusters. **(B)** Example of TADpole tool applied to a 10Mb-region from a Hi-C matrix of human chr18 at 40kb resolution. The complete dendrogram (*left*) is cut using the broken-stick model to prune nonsignificant partitions. The first (2 clusters) and last (21 clusters) significant subdivisions are shown as light blue dashed lines. A selection of the corresponding partitions (*right*) is mapped on the analyses Hi-C matrix as light grey dashed lines. The optimal division in 16 clusters, identified by the highest Calinski-Harabasz (CH) index, is shown as a dark blue dashed line.

#### (i) Preprocessing of the input dataset

TADpole is designed to process all-vs-all intra-chromosomal interactions matrices representing an entire chromosome, or a continuous chromosome region. Input data are formatted as matrices containing the interaction values in each cell *ij*. An optional filtering step can be applied to exclude columns with a low number of interactions, which are typically local biases (42). Specifically, the rows (and columns) that contain an empty cell at the main diagonal, or those whose cumulative interactions are below the first percentile (default) are excluded from the analysis. To enhance the signal-to-noise ratio, the interaction matrix is transformed into its Pearson correlation coefficient (PCC) matrix as in (13), and a principal component analysis (PCA) is performed using the *prcomp* function from the *stats* R package (R Core Team, 2013). Only the first *N_PC_* (by default, 200) principal components are retained, which are enough to extract more than 85% of the variance in the test datasets (**Supplementary Figure 1**). To reduce memory usage and processing time, TADpole has the option to divide the interaction matrix by the centromere (considered to be the longest contiguous stretch of columns with no interactions in the Hi-C matrix) and process each chromosomal arm separately. This is particularly recommended when working with matrices of more than 15,000 bins.

#### (ii) Constrained hierarchical clustering optimization

Per each value of *N_PC_*, the dimensionally-reduced matrix is transformed into a Euclidean distance matrix. This distance matrix is then partitioned into topological domains using a constrained hierarchical clustering procedure as implemented in the Constrained Incremental Sums of Squares clustering method (*coniss*) of the *rioja* R package (Juggins *et al*., 2017). This analysis explicitly assumes the following two priors: first, the genome is organized in a hierarchical manner, with higher-order structures containing lower-order ones, and second, every pair of contiguous genomic loci must either belong to the same self-interacting domain or to the immediately contiguous one. The constrained hierarchical clustering results in a tree-like description of the organization of the genome. Next, using the broken-stick model as implemented in the *bstick* function from the *rioja* R package (Juggins *et al*., 2017), the dendrogram is cut at the sensible maximum number of statistically significant clusters (*max*(N_D_)). Next, the Calinski-Harabasz (CH) index is computed per each of the obtained significant partitions using the *calinhara* function from the *fpc* R package (Henning C, 2018). The maximum CH is associated to the optimal chromatin subdivision, while all the other significant hierarchical levels correspond the ones with the optimal number of first *N_PC_* (**Figure 1B**).

#### (iii) Genome Segmentation

TADpole generates four main descriptors that recapitulate the entire sets of results, namely: (i) the optimal number of principal components; (ii) the dendrogram cut at the maximum significant level; (iii) the start and end coordinates of the domains and the CH index per each significant level; (iv) and the optimal number of domains. All the TADpole output is organized in a comprehensive R object.

### TADpole benchmark analysis

#### Benchmark Hi-C dataset and scripts

A pre-existing benchmark dataset, that comprises Hi-C interaction matrices of the entire chromosome 6 in the human cell line GM12787, was used for the analysis (29). A total of 24 different conditions were tested: (i) twelve matrices given by the combination of four different resolutions (10kb, 50kb, 100kb and 250kb) and three normalization strategies (raw, Iterative Correction and Eigenvector decomposition (ICE) (14) and parametric model of Local Genomic Feature (LGF) (43), and (ii) twelve matrices obtained by downsampling the ICE interaction matrix at 50kb resolution (**Figure 2A**). The scripts for benchmarking were downloaded and used as released in the repository https://github.com/CSOgroup/TAD-benchmarking-scripts (29) (**Data availability**). The processed Hi-C dataset was shared by Zufferey and colleagues, this eliminated from the analysis possible biases associated with the use of different pipelines for Hi-C interaction data reconstruction (28). To compare on equal footing with the other 22 TAD callers analyzed using the same benchmark, only levels of division that comprise at least 10 chromatin domains were taken into consideration for the analysis. Within these levels, the optimal partition was identified using the CH index as described before.

**Figure 2.**
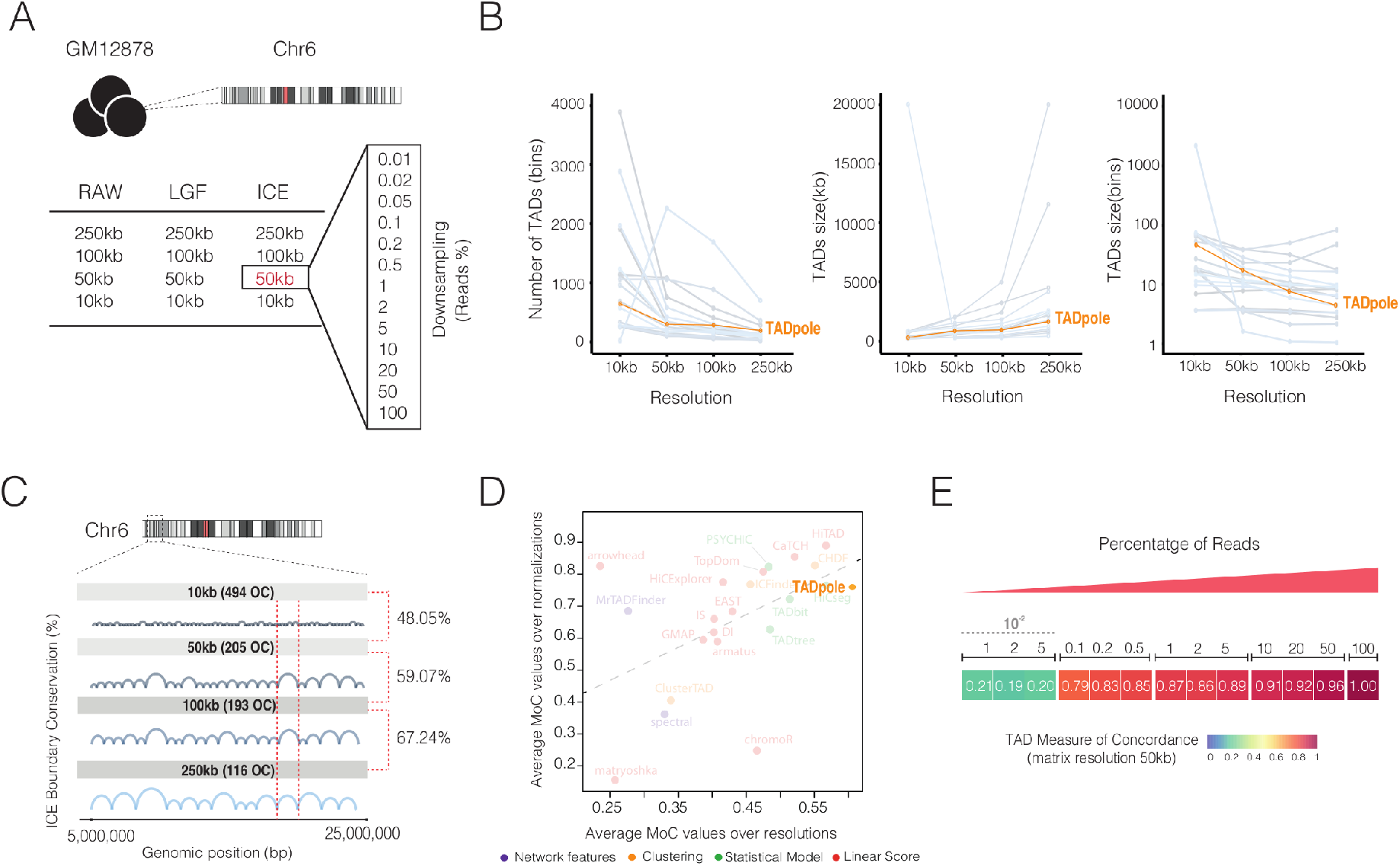
Technical benchmark of TADpole tool. **(A)** Descriptions of the dataset used for the TADpole benchmark analysis of Zufferey *et al*. (29). The dataset includes the Hi-C interactions matrices for chromosome 6 in GM12878 cells organized in 24 different forms (**Material and Methods**). **(B-E)** Technical benchmark of TADpole optimal partition and comparison with 22 other TAD callers considered in (29). **(B)** Analysis of the number of TADs at different matrix binning and the TAD size in terms of kb and number of bins. **(C)** Fraction of conserved TADs boundaries at different resolutions in the entire chromosome 6. The scheme illustrates the analysis focusing on the region from 5 to 25Mb. **(D)** Average Measure of Concordance (MoC) values across normalizations *vs* the average MoC value across resolutions. The color scheme reflects the specific approach used in each TAD caller. **(E)** The MoC values over different downsampling levels of the ICE-normalized interaction matrix at 50kb. Panel B and D have been adapted from Figure 2C of Zufferey *et al*. (29) to include TADpole in the comparison of TAD callers.

#### The technical benchmark

TADpole optimal topological partitions were compared over different resolutions, normalization strategies, and sequencing depths as previously described (29). To compare the conservation of the TAD borders between two partitions, two different metrics were applied:

1. The overlap score (29) was used to compare partitions across different resolutions. This is the percentage of overlapping borders, with one bin of tolerance. The statistical significance of each overlap score was estimated by drawing 10,000 random partitions at the finer resolution (preserving the number of optimal clusters of the real case) and computing their overlap with the subdivision at the coarser resolution. The p-value of the real-case overlap was computed as the fraction of randomized partitions with larger overlap.
2. The Measure of Concordance (MoC) (29), was used to compare partitions across different resolutions and normalization strategies. MoC ranges from 1 for a perfect match to 0 for poorly scoring comparisons.

#### The biological benchmark

To test the biological relevance of the TADs identified by TADpole, the enrichment of main architectural proteins determined at TAD borders (CTCF, SMC3, and RAD21) and within TADs (H3K27me3, and H3K36me3) were studied. CTCF and cohesin sub-units, such as SMC3, and RAD21 have been shown, in fact, to be enriched at TAD borders (15,44) while H3K27me3 and H3K36me3 marks have been related to acting as a differentiator of TADs because topological domains are enriched in either one or the other, but not both (13,15,20,21). The ChIP-seq profiles were downloaded from ENCODE (45) (https://www.encodeproject.org/) (**Supplementary Table 1)**. For each protein, a *consensus* profile was determined as the intersection of the peaks identified in each experiment using the *multiIntersectBed* function from the BEDTools suite (Quinlan and Hall, 2010). Similar to Zufferey *et al*. (29), the fold change enrichments of CTCF, RAD21 and SMC3 at TAD borders, and the H3k27me3/H3k36me3 log10-ratio for a given partition were computed.

### Difference score between topological partitions (DiffT)

The TADpole tool was next applied to two Capture Hi-C (cHi-C) datasets in embryonic day E11.5 mouse limb buds (41). Specifically, two homozygous strains were considered comprising the wild type (WT) and the so-called inversion1 (Inv1). The cHi-C interaction maps were downloaded from GEO (46) at the GSM3261968 and GSM3261969 entries for WT and Inv1, respectively. The region chr1:73.92-75.86 Mb was extracted and used for further analysis.

To compare the WT and Inv1 partitions identified by TADpole at a fixed level of the hierarchy, we defined a difference topology score (DiffT). Specifically, the partitioned matrices were transformed into binary forms *W* for WT, and analogously *V* for Inv1, in which each entry *w_ij_* (*v_ij_*) is equal to 1 if the bins *i* and *j* are in the same TAD and 0 otherwise. Then, DiffT is computed as the normalized (from 0 to 1) difference between the binarized matrices as a function of the bin index *l* as:

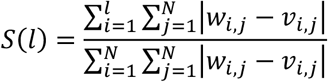

where *N* is the size of the matrix.

To test whether the identified partition in Inv1 is different from WT, at each level of the chromatin hierarchy, a statistical analysis was introduced. This analysis assesses the significance of DiffT at each bin of the matrix. A total of 10,000 random partitions of the region were simulated excluding the bad columns of the Inv1 matrix. The DiffT score was computed between simulated and WT partitions (DiffT_simulated-wt_). At each bin, the fraction of DiffT_simulated-wt_ lower or equal to the DiffT_Inv1-wt_ score estimates the *p-value*. A p-value < 0.05 means that a significant difference is located from the bin under consideration onwards. Hence, the bin with the minimum p-value marks the starting point of the genomic region where the most significant fraction of the DiffT score is located.

## RESULTS

### TADpole benchmark analysis

To quantitatively compare TADpole with other 22 TAD callers, we applied the multiple conditions test proposed in (29) on the same reference benchmark dataset (**Figure 2A** and **Material and Methods**).

#### Technical benchmarking

We assessed various technical aspects of TADpole as well as the robustness of TADpole identified domains with respect to different resolutions, normalization strategies, and sequencing depths of the input matrix (**Figure 2A**). Firstly, we examined the number and the size (in kilobases and in bins) of the optimal domain partition of the ICE normalized maps at different resolutions (**Figure 2B**). We found that, as the resolution of the Hi-C interaction map decreased, both the numbers of TADs and the mean TAD size in bins decreased with a 4-fold reduction. TADpole followed a similar grow tendency (positive when the TADs are measure in kilobases and negative with the TADs are measured in bins) as the majority of the other TAD callers independently on the normalized strategy applied (**Supplementary Table 2**).

We also inspected if TADpole identified robust boundaries that were conserved at different resolutions. To measure this conservation, we tested if a border detected in the ICE normalized Hi-C matrices at a certain resolution was conserved in the resolution immediately finer (**Figure 2C**). At the coarser resolutions, that is 250kb *vs*. 100kb, we found a high agreement (67%), that decreased only slightly to (59%) at intermediate ones (100 *vs*. 50kb). Interestingly, we found that even at the finer resolutions (50kb *vs*. 10kb), where the 48% of the borders were conserved, this analysis was consistent with a statistically significant overlap (p-value<0.05).

Next, we used the Measure of Concordance (MoC) (**Material and Methods**) to estimate if the number and the position of the borders of chromatin domains identified by TADpole were affected by the matrix resolutions and by different normalization strategies. Interestingly, we found that the MoC over different matrix resolutions had values in the [0.45:0.82] range with an average MoC of 0.63, and ranked first when compared with the other 22 TAD callers previously benchmarked (29). TADpole was also robust over different normalization strategies with an average MoC of 0.74, ranking 9^th^ over the 22 TAD callers. Comparing the average of resolutions *vs*. normalizations MoC values of TADpole with the rest of TADcallers (**Figure 2D**), we found that TADpole appeared in the top-right corner of the plot demonstrating its overall high robustness and confidence to identify optimal chromatin domains independently of the resolution or the normalization of the input Hi-C matrix.

We also tested the TADpole propensity to identify consistent optimal chromatin domains independently of the sequencing depth (**Figure 2E**). We compared the partitions obtained by doing 12 different sub-sampling of the ICE-normalized interaction matrix at 50kb with the full interaction matrix using the MoC. We found that TADpole partitions were clearly robust to down-sampling with a MoC score of 0.79 with just 0.1% of the total data. This feature classified TADpole as the top TAD caller with respect to the other 22 tools.

#### Biological benchmarking

With the lack of a gold standard to define TADs in Hi-C interaction maps (28,29), we investigated the biological relevance of the domains identified by TADpole in terms of their association with biological features that have been shown to have an important role in the formation and maintenance of TADs. We found that the intensity profiles of the CTCF, RAD21, and SMC3 signals were peaked at TADpole chromatin borders (**Figure 3A**). To compare these results with the set of other TAD callers, we computed the fold change enrichments at the peak with respect to the flanking regions (**Figure 3B**). TADpole resulted in a fold change enrichment around 1 for each of the three architectural proteins (1.18 in CTCF, 1.06 in RAD21 and 0.97 in SMC3, respectively), that was consistent with a significantly high peak at the border compared with the background (p-value<10^−5^). In this analysis, TADpole ranked as the 6^th^ TAD caller. Additionally, more than 40% of the tagged boundaries are enriched in one or more of these three architectural proteins being CTCF (42%) and SMC3 (42%) the most abundant ones. Considering the enrichment at TADs boundaries, TADpole ranked 3^rd^ within the set of 22 TAD examined callers (**Figure 3C**). To further study the enrichment of these biological features at domain borders, we performed an analysis of the fold change of CTCF, RAD21 and SMC3 in each of the chromosome arms. In all the identified levels, there was a positive fold-change with certain variability with the level of nested data, being CTCF in all the cases, the most enriched architectural protein in the border regions (**Figure 3D**).

**Figure 3.**
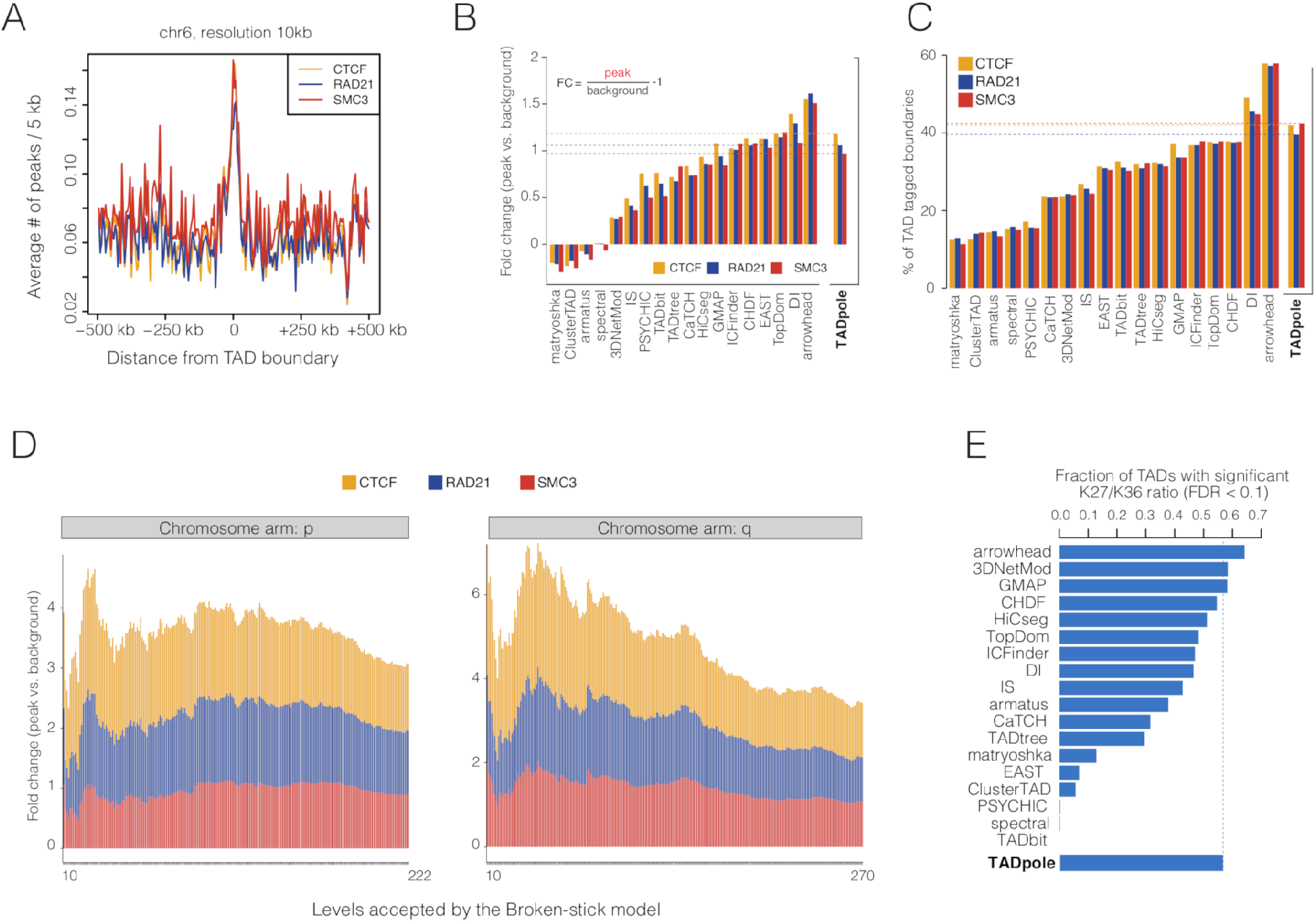
Biological benchmark of TADpole tool. TADpole borders for the optimal level of the hierarchical partition and chromatin-organizer proteins. **(A)** The mean ChIP-seq signal profiles (5-kb intervals) in 1Mb region around the TADpole domains borders are shown for CTCF, RAD21 and SMC3. **(B)** The fold-change enrichment of CTCF, RAD21 and SMC3 at domain borders *vs*. background and **(C)** the percentage identified TADs boundaries tagged with CTCF, RAD21 and SMC3 are shown as bar plots. **(D)** Cumulative fold-change of the enrichment in CTCF, RAD21 and SMC3 at domain borders *vs*. background for all the significant levels (minimum 10 partitions as in (29)) retrieved by the broken-stick model in the two chromosomes arms (p, q) of chromosome 6. **(E)** The fractions of TADs with significant log10 ratio between H3K36me3 and H3K27me3 (**Material and Methods**) in TADpole and the 22 TAD callers are represented as bar plots. Panels B, C and E have been adapted from Figure 5D, 5E, 5I of Ref. (29) reporting on the results of the other TAD callers.

TADs are usually expected to be transcriptionally either active or inactive (15) with the TAD body enriched in active or inactive histone mark. To assess if the interior of TADpole detected partitions was indeed enriched in either active or inactive chromatin, we considered the signals of two marks H3K36me3 for transcriptional activity and H3K27me3 for repression, and measured the fraction of TADs where the Log10 of their ratio (H3K27me3/H3K36me3) was significantly high or low (FDR<0.1). Notably, we found that the majority (57%) of the TADpole identified TADs have a defined active or inactive state, locating TADpole within the top four TAD callers based on this criterion (**Figure 3E**).

### Applications to capture Hi-C datasets

To show the effectiveness of TADpole, we applied our caller to cHi-C experiment of embryonic day E11.5 mouse cells (41). Kraft *et al*. investigated the pathogenic consequences of balanced chromosomal rearrangements in embryonic mouse limb buds, focusing on a 1.9Mb region in chr1 (chr1:73.92-75.86 Mb) where a cluster of genomic regulators of *Epha4 locus* is located. The authors generated, together with the wild-type (WT), a total of 4 mutant strains, each inducing different inversions. We compared the WT strain with the sole inversion producing a homozygous strain (here called Inv1), that is located between the telomeric site of *Epha4* enhancer cluster and the promoters of *Resp18* (breakpoint at chr1:75,275,966-75,898,706 **Figure 4A**). Analysis of the entire TADpole dendrogram revealed the existence of 19 and 17 significant partition levels in WT and Inv1, respectively. The optimal ones were 11 for WT and 2 for Inv1. At a visual inspection, the maps in **Figure 4A** show that the difference between the chromatin partitions increases with the partition level, and accumulates in the region of the inversion.

**Figure 4.**
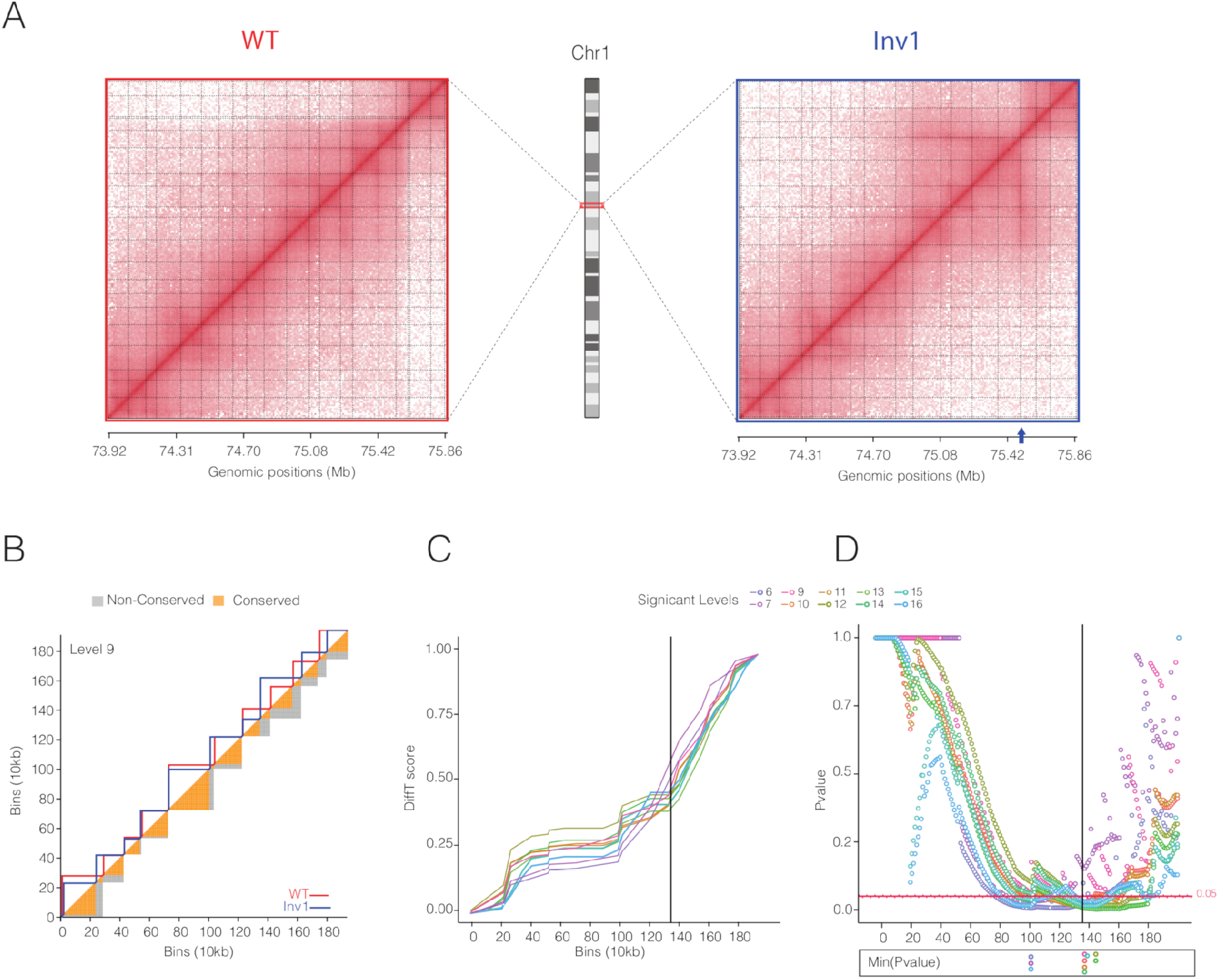
Characterization of topological difference in capture Hi-C datasets. **(A)** Capture Hi-C maps of chr1:73.92-75.86Mb region in wild-type (WT) and inversion 1 (Inv1) strains (41). In both matrices, TADpole significant partitions are shown as light gray dashed lines. The blue arrow indicates the centromeric breakpoint located at the promoter of *Resp18* gene in Inv1. **(B)** DiffT score scheme for level 9. The upper triangle of the matrix shows the TADs borders identified by TADpole in WT and Inv1 matrices as red and blue continuous lines, respectively. The lower triangle of the matrix shows the conserved and non-conserved areas of the TADs in orange and gray, respectively. In the panels C and D, the Inv1 breakpoint is highlighted with a solid black line and only the significant levels (with at least a p-value < 0.05) are shown. **(C)** DiffT score profiles as a function of the matrix bins (**Material and Methods**). **(D)** P-value profiles per bin for automated detection of significant differences. In the lower panel, the bins associated with minimum p-values per level are marked with empty dots.

To statistically quantify and localize the significant topological differences between the WT and Inv1 matrices, we computed their DiffT score profiles (**Material and Methods** and **Figure 4B** for the partition in 9 domains), at each level of the hierarchical partition. We found that the DiffT profiles sharply increased close to the point of the inversion (**Figure 4C**). Based on the p-value profiles (**Figure 4D**), we identified two regions where the minimum p-values (one per partition level) accumulated. Notably, 70% of minimum p-values were located within a region, spanning 50kb, from the point where inversion was induced, suggesting that the significant topological changes between WT and Inv1 accumulated in the inverted region.

## DISCUSSION AND CONCLUSION

In this work we introduced TADpole, a tool to identify hierarchical topological domains from all-vs-all intra-chromosomal interaction matrices. In line with previously introduced concepts such as metaTADs (26) and sub-TADs (27), we propose that there is not a single meaningful subdivision of chromatin domains, but rather a set of hierarchical levels associated with different genomic features. Notably, TADpole characterizes the entire hierarchy of TADs while assessing the significance at various levels of partition, paving the way for deepening our understanding of the nested topology and its biological role. Indeed, the principles behind this nested structure are not yet fully understood. Different levels of the hierarchy can be involved in the dynamical modulations of TADs upon perturbation, responding to or causing changes in gene regulation (3,22,47). Alternatively, these nested organization can rise as effect of the variability in the topological conformation observed in individual cells (48). However, we cannot exclude the possibility that this nested structure exists in single cells as multi-site interactions conformation acting together to establish robust gene regulation networks. All these evidences highlight the importance to have a tool like TADpole, that systematically characterizes the entire TAD architecture from all-vs-all intra-chromosomal interaction matrices.

Here, we compared TADpole’s performance with a set of other 22 TAD callers following the benchmark analysis performed by Zufferey *et al*. (29). TADpole identifies a number of TADs over different resolutions that is in agreement with other TAD callers (**Figure 2B**). The identified domains have an average size of 855kb, in agreement with the reported average TADs size in mammalian cells (~900-1000kb) (15). TADpole shows one of the largest consistencies over different normalization strategies (including also non-normalized data), resolutions and sequencing depths (**Figure 2D** and **Supplementary Table 2**). These observations make TADpole potentially suitable for analyzing sparse datasets. TADpole has been shown to be technically robust. Indeed, the optimal TADpole chromatin partition borders present a high enrichment of architectural proteins such as CTCF, SMC3, and RAD21, and this enrichment is maintained, for certain partitions, over all the significant hierarchical levels (**Figure 3D**). The identified TADs are enriched in either active (H3K36me3) or inactive (H3K27me3) marks (**Figure 3E**), suggesting a strong consistency between the structural definition of TADs and their biological characterization. A handful of algorithms for the detection of chromatin domains as nested TADs have been implemented (CaTCH (36), GMAP (37), matryoshka (38), and PSYCHIC (39)). However, the uniqueness of TADpole is its ability to provide multiple significant partitions and define the optimal one in an unsupervised manner by using the Broken-Stick model and the Calinski-Harabasz index criteria. Notably, in the benchmark analysis presented here (**Figures 2** and **3**), TADpole performs generally better than all the other nested TAD callers when considering the technical and biological benchmarking performed here.

A possible advantage of TADpole over existing TAD callers is the pre-processing data step. Indeed, the PCC transformation and the PCA application regularize the input matrix so that the specific normalization applied on the input and the sparsity of the data have little effect on identifying TADs. Previously, other architectural features of the chromatin have been already studied using PCA. The first principal component is widely used to identify the chromatin segregation into compartments (13). The second and the third PCs have been associated instead to intra-arm features mainly centromere-centromere and telomere-telomere interactions enrichment (14). Moreover, the first PCs have been used to assess the similarity between two interaction maps (14) as well as to quantify their reproducibility (49). Here we have demonstrated that there exists an optimal set of PCs capable of identifying the hierarchical structure of chromatin, extending the current application of PCA to characterize genome topology.

We provide a proof of TADpole’s usability on a topologically complex region analyzing cHi-C data in both a wild-type strain and a mutant one carrying a genomic inversion (41) (**Figure 4B**). The use of TADpole in combination with the DiffT score is able to identify the inverted region as the one with the highest difference in topological partitions, proving that this strategy can isolate bin-dependent and statistically significant topological dissimilarities. Overall, we prove that the DiffT score allows to evaluate *a priori* the location where the most significant topological differences between two hierarchical subdivisions are accumulated.

In summary, TADpole combines straightforward bioinformatic analyses such as PCA and hierarchical clustering to study continuous nested hierarchical segmentation of an all-vs-all intra-chromosomal interactions matrix. Additionally, we demonstrated the technical and biological robustness of TADpole, and its usability in identifying topological difference in high-resolution capture Hi-C experiments. TADpole is released as a publicly-available, open-source and numerically-efficient R tool. As such, TADpole represents a comprehensive tool that fulfils the needs of the scientific community for an accurate TAD caller able to comprehensively study the interplay between the hierarchical chromatin topology and genomic function.

## DATA AVAILABILITY

The TADpole is freely available for download as an R package at https://github.com/3DGenomes/TADpole. The scripts for the technical and biological benchmarks were obtained from the repository https://github.com/CSOgroup/TAD-benchmarking-scripts (28). Specifically, the script *fig2_fig3_fig4_fig5_moc_calc.R* was used for panels **Figure 2B** to **E**, the script *StructProt_EnrichBoundaries_script.R* for panels **Figure 3A** to **D**, and the script *HistMod_script.sh* for panel in **Figure 3E**. Default parameters were applied.

## ACKNOWLEDGEMENT

We thank Dr. M. Zufferey and Dr. G. Ciriello for providing us with the dataset used in Ref. (29), that make possible an easy and quick comparison of TADpole with 22 other TAD callers. We thank also Dr. K. Kraft and Dr. S. Mundlos for helping in the interpretation of the Capture Hi-C datasets in Ref. (41). We acknowledge the ENCODE consortium and the ENCODE production laboratories that generated the datasets used in the manuscript.

---

**Supplementary Figure 1.**
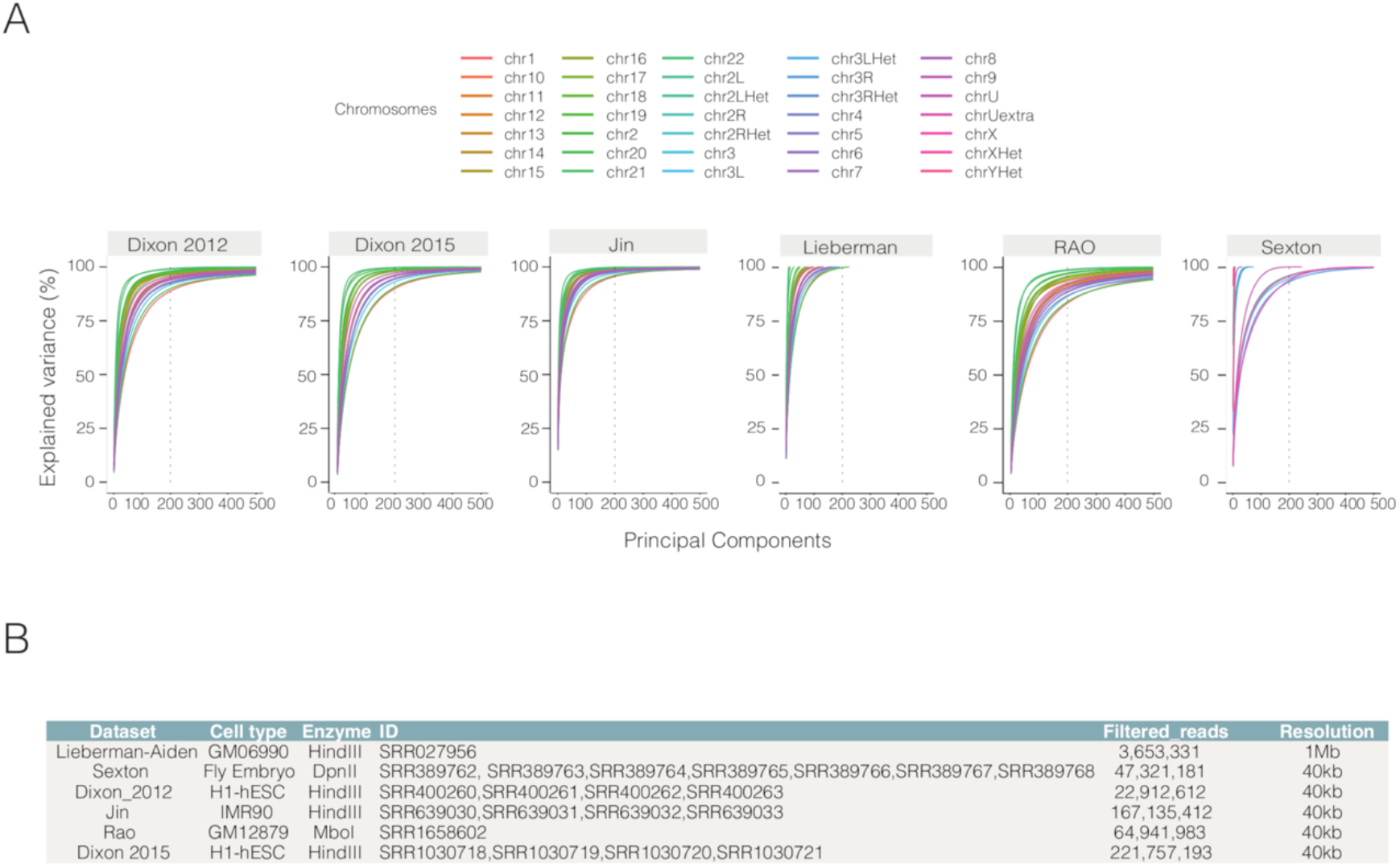
Percentage of explained variance as a function of the number of retained principal components of for various datasets. **(A)** Each continuous line represents a different chromosome, and the vertical dashed lines mark the default maximum value of (200) first PCs retained in TADpole. **(B)** The Hi-C experimental datasets used characterized with five descriptors: cell type, restriction enzyme, the NCBI accession numbers, number of the valid reads retrieved after filtering using an in-house pipeline based on TADbit (50), and binning. The datasets from different NCBI entries are merged and the resulting matrices after filtering are binned using an equal bin-width of 40kb, but for Lieberman-Aiden dataset (13) in which bin-width is 1Mb.

**Supplementary Table 1.** Encode IDs of the Chip-seq experiments used in the biological benchmark analysis.

| Chromatin Marks | Encode ID |
| --- | --- |
| CTCF | ENCSR000DRZ, ENCSR000DKV, ENCSR000DZN, ENCSR000AKB |
| SMC3 | ENCSR000DZP |
| RAD21 | ENCSR000BMY, ENCSR000EAC |
| H3K36me3 | ENCSR000DRW |
| H3K27me3 | ENCSR000DRX |

**Supplementary Table 2.** The total number of TADs and the corresponding average size detected in raw and normalized (by ICE and LGF) Hi-C matrices across different resolutions.

| Resolution | Raw<br>N° TADs | Raw<br>Size (kb) | Raw<br>Size(bins) | ICE<br>N° TADs | ICE<br>Size (kb) | ICE<br>Size (bins) | LGF<br>N° TADs | LGF<br>Size (kb) | LGF<br>Size (bins) |
| --- | --- | --- | --- | --- | --- | --- | --- | --- | --- |
| 250kb | 118 | 1425.85 | 5.7 | 116 | 1465.52 | 5.86 | 120 | 1402.08 | 5.61 |
| 100kb | 193 | 868.91 | 8.68 | 193 | 879.79 | 8.79 | 193 | 884.46 | 88.45 |
| 50kb | 217 | 772.35 | 15.45 | 205 | 814.39 | 16.29 | 208 | 805.77 | 16.12 |
| 10kb | 535 | 313.01 | 31.3 | 494 | 338.7 | 33.87 | 510 | 328.29 | 32.83 |

## REFERENCES

1. Sexton, T. and Cavalli, G. (2015) The role of chromosome domains in shaping the functional genome. Cell, 160, 1049–1059.

2. Dekker, J. and Mirny, L. (2016) The 3D Genome as Moderator of Chromosomal Communication. Cell, 164, 1110–1121.

3. Stadhouders, R., Vidal, E., Serra, F., Di Stefano, B., Le Dily, F., Quilez, J., Gomez, A., Collombet, S., Berenguer, C., Cuartero, Y. et al. (2018) Transcription factors orchestrate dynamic interplay between genome topology and gene regulation during cell reprogramming. Nat Genet, 50, 238–249.

4. Paulsen, J., Liyakat Ali, T.M., Nekrasov, M., Delbarre, E., Baudement, M.O., Kurscheid, S., Tremethick, D. and Collas, P. (2019) Long-range interactions between topologically associating domains shape the four-dimensional genome during differentiation. Nat Genet, 51, 835–843.

5. Bonev, B., Mendelson Cohen, N., Szabo, Q., Fritsch, L., Papadopoulos, G.L., Lubling, Y., Xu, X., Lv, X., Hugnot, J.P., Tanay, A. et al. (2017) Multiscale 3D Genome Rewiring during Mouse Neural Development. Cell, 171, 557–572 e524.

6. Zheng, H. and Xie, W. (2019) The role of 3D genome organization in development and cell differentiation. Nat Rev Mol Cell Biol.

7. Lupianez, D.G., Kraft, K., Heinrich, V., Krawitz, P., Brancati, F., Klopocki, E., Horn, D., Kayserili, H., Opitz, J.M., Laxova, R. et al. (2015) Disruptions of topological chromatin domains cause pathogenic rewiring of gene-enhancer interactions. Cell, 161, 1012–1025.

8. Franke, M., Ibrahim, D.M., Andrey, G., Schwarzer, W., Heinrich, V., Schopflin, R., Kraft, K., Kempfer, R., Jerkovic, I., Chan, W.L. et al. (2016) Formation of new chromatin domains determines pathogenicity of genomic duplications. Nature, 538, 265–269.

9. Groschel, S., Sanders, M.A., Hoogenboezem, R., de Wit, E., Bouwman, B.A.M., Erpelinck, C., van der Velden, V.H.J., Havermans, M., Avellino, R., van Lom, K. et al. (2014) A single oncogenic enhancer rearrangement causes concomitant EVI1 and GATA2 deregulation in leukemia. Cell, 157, 369–381.

10. Flavahan, W.A., Drier, Y., Liau, B.B., Gillespie, S.M., Venteicher, A.S., Stemmer-Rachamimov, A.O., Suva, M.L. and Bernstein, B.E. (2016) Insulator dysfunction and oncogene activation in IDH mutant gliomas. Nature, 529, 110–114.

11. Krijger, P.H. and de Laat, W. (2016) Regulation of disease-associated gene expression in the 3D genome. Nat Rev Mol Cell Biol, 17, 771–782.

12. Cremer, T. and Cremer, C. (2001) Chromosome territories, nuclear architecture and gene regulation in mammalian cells. Nat Rev Genet, 2, 292–301.

13. Lieberman-Aiden, E., van Berkum, N.L., Williams, L., Imakaev, M., Ragoczy, T., Telling, A., Amit, I., Lajoie, B.R., Sabo, P.J., Dorschner, M.O. et al. (2009) Comprehensive mapping of long-range interactions reveals folding principles of the human genome. Science, 326, 289–293.

14. Imakaev, M., Fudenberg, G., McCord, R.P., Naumova, N., Goloborodko, A., Lajoie, B.R., Dekker, J. and Mirny, L.A. (2012) Iterative correction of Hi-C data reveals hallmarks of chromosome organization. Nat Methods, 9, 999–1003.

15. Rao, S.S., Huntley, M.H., Durand, N.C., Stamenova, E.K., Bochkov, I.D., Robinson, J.T., Sanborn, A.L., Machol, I., Omer, A.D., Lander, E.S. et al. (2014) A 3D map of the human genome at kilobase resolution reveals principles of chromatin looping. Cell, 159, 1665–1680.

16. Nir, G., Farabella, I., Perez Estrada, C., Ebeling, C.G., Beliveau, B.J., Sasaki, H.M., Lee, S.D., Nguyen, S.C., McCole, R.B., Chattoraj, S. et al. (2018) Walking along chromosomes with super-resolution imaging, contact maps, and integrative modeling. PLoS Genet, 14, e1007872.

17. Boettiger, A.N., Bintu, B., Moffitt, J.R., Wang, S., Beliveau, B.J., Fudenberg, G., Imakaev, M., Mirny, L.A., Wu, C.T. and Zhuang, X. (2016) Super-resolution imaging reveals distinct chromatin folding for different epigenetic states. Nature, 529, 418–422.

18. Bintu, B., Mateo, L.J., Su, J.H., Sinnott-Armstrong, N.A., Parker, M., Kinrot, S., Yamaya, K., Boettiger, A.N. and Zhuang, X. (2018) Super-resolution chromatin tracing reveals domains and cooperative interactions in single cells. Science, 362.

19. Szabo, Q., Jost, D., Chang, J.M., Cattoni, D.I., Papadopoulos, G.L., Bonev, B., Sexton, T., Gurgo, J., Jacquier, C., Nollmann, M. et al. (2018) TADs are 3D structural units of higher-order chromosome organization in Drosophila. Sci Adv, 4, eaar8082.

20. Dixon, J.R., Selvaraj, S., Yue, F., Kim, A., Li, Y., Shen, Y., Hu, M., Liu, J.S. and Ren, B. (2012) Topological domains in mammalian genomes identified by analysis of chromatin interactions. Nature, 485, 376–380.

21. Nora, E.P., Lajoie, B.R., Schulz, E.G., Giorgetti, L., Okamoto, I., Servant, N., Piolot, T., van Berkum, N.L., Meisig, J., Sedat, J. et al. (2012) Spatial partitioning of the regulatory landscape of the X-inactivation centre. Nature, 485, 381–385.

22. Le Dily, F., Baù, D., Pohl, A., Vicent, G.P., Serra, F., Soronellas, D., Castellano, G., Wright, R.H., Ballare, C., Filion, G. et al. (2014) Distinct structural transitions of chromatin topological domains correlate with coordinated hormone-induced gene regulation. Genes Dev, 28, 2151–2162.

23. Le Dily, F., Vidal, E., Cuartero, Y., Quilez, J., Nacht, A.S., Vicent, G.P., Carbonell-Caballero, J., Sharma, P., Villanueva-Canas, J.L., Ferrari, R. et al. (2019) Hormone-control regions mediate steroid receptor-dependent genome organization. Genome Res, 29, 29–39.

24. Dixon, J.R., Jung, I., Selvaraj, S., Shen, Y., Antosiewicz-Bourget, J.E., Lee, A.Y., Ye, Z., Kim, A., Rajagopal, N., Xie, W. et al. (2015) Chromatin architecture reorganization during stem cell differentiation. Nature, 518, 331–336.

25. Bonev, B. and Cavalli, G. (2016) Organization and function of the 3D genome. Nat Rev Genet, 17, 661–678.

26. Fraser, J., Ferrai, C., Chiariello, A.M., Schueler, M., Rito, T., Laudanno, G., Barbieri, M., Moore, B.L., Kraemer, D.C., Aitken, S. et al. (2015) Hierarchical folding and reorganization of chromosomes are linked to transcriptional changes in cellular differentiation. Mol Syst Biol, 11, 852.

27. Berlivet, S., Paquette, D., Dumouchel, A., Langlais, D., Dostie, J. and Kmita, M. (2013) Clustering of tissue-specific sub-TADs accompanies the regulation of HoxA genes in developing limbs. PLoS Genet, 9, e1004018.

28. Forcato, M., Nicoletti, C., Pal, K., Livi, C.M., Ferrari, F. and Bicciato, S. (2017) Comparison of computational methods for Hi-C data analysis. Nat Methods, 14, 679–685.

29. Zufferey, M., Tavernari, D., Oricchio, E. and Ciriello, G. (2018) Comparison of computational methods for the identification of topologically associating domains. Genome Biol, 19, 217.

30. Crane, E., Bian, Q., McCord, R.P., Lajoie, B.R., Wheeler, B.S., Ralston, E.J., Uzawa, S., Dekker, J. and Meyer, B.J. (2015) Condensin-driven remodelling of X chromosome topology during dosage compensation. Nature, 523, 240–244.

31. Oluwadare, O. and Cheng, J. (2017) ClusterTAD: an unsupervised machine learning approach to detecting topologically associated domains of chromosomes from Hi-C data. BMC Bioinformatics, 18, 480.

32. Haddad, N., Vaillant, C. and Jost, D. (2017) IC-Finder: inferring robustly the hierarchical organization of chromatin folding. Nucleic Acids Res, 45, e81.

33. Filippova, D., Patro, R., Duggal, G. and Kingsford, C. (2014) Identification of alternative topological domains in chromatin. Algorithms Mol Biol, 9, 14.

34. Weinreb, C. and Raphael, B.J. (2016) Identification of hierarchical chromatin domains. Bioinformatics, 32, 1601–1609.

35. Norton, H.K., Emerson, D.J., Huang, H., Kim, J., Titus, K.R., Gu, S., Bassett, D.S. and Phillips-Cremins, J.E. (2018) Detecting hierarchical genome folding with network modularity. Nat Methods, 15, 119–122.

36. Zhan, Y., Mariani, L., Barozzi, I., Schulz, E.G., Bluthgen, N., Stadler, M., Tiana, G. and Giorgetti, L. (2017) Reciprocal insulation analysis of Hi-C data shows that TADs represent a functionally but not structurally privileged scale in the hierarchical folding of chromosomes. Genome Res, 27, 479–490.

37. Yu, W., He, B. and Tan, K. (2017) Identifying topologically associating domains and subdomains by Gaussian Mixture model And Proportion test. Nat Commun, 8, 535.

38. Malik, L. and Patro, R. (2018) Rich Chromatin Structure Prediction from Hi-C Data. IEEE/ACM Trans Comput Biol Bioinform.

39. Ron, G., Globerson, Y., Moran, D. and Kaplan, T. (2017) Promoter-enhancer interactions identified from Hi-C data using probabilistic models and hierarchical topological domains. Nat Commun, 8, 2237.

40. Dryden, N.H., Broome, L.R., Dudbridge, F., Johnson, N., Orr, N., Schoenfelder, S., Nagano, T., Andrews, S., Wingett, S., Kozarewa, I. et al. (2014) Unbiased analysis of potential targets of breast cancer susceptibility loci by Capture Hi-C. Genome Res, 24, 1854–1868.

41. Kraft, K., Magg, A., Heinrich, V., Riemenschneider, C., Schopflin, R., Markowski, J., Ibrahim, D.M., Acuna-Hidalgo, R., Despang, A., Andrey, G. et al. (2019) Serial genomic inversions induce tissue-specific architectural stripes, gene misexpression and congenital malformations. Nat Cell Biol, 21, 305–310.

42. Vidal, E., le Dily, F., Quilez, J., Stadhouders, R., Cuartero, Y., Graf, T., Marti-Renom, M.A., Beato, M. and Filion, G.J. (2018) OneD: increasing reproducibility of Hi-C samples with abnormal karyotypes. Nucleic Acids Res, 46, e49.

43. Hu, M., Deng, K., Selvaraj, S., Qin, Z., Ren, B. and Liu, J.S. (2012) HiCNorm: removing biases in Hi-C data via Poisson regression. Bioinformatics, 28, 3131–3133.

44. Kojic, A., Cuadrado, A., De Koninck, M., Gimenez-Llorente, D., Rodriguez-Corsino, M., Gomez-Lopez, G., Le Dily, F., Marti-Renom, M.A. and Losada, A. (2018) Distinct roles of cohesin-SA1 and cohesin-SA2 in 3D chromosome organization. Nat Struct Mol Biol, 25, 496–504.

45. Davis, C.A., Hitz, B.C., Sloan, C.A., Chan, E.T., Davidson, J.M., Gabdank, I., Hilton, J.A., Jain, K., Baymuradov, U.K., Narayanan, A.K. et al. (2018) The Encyclopedia of DNA elements (ENCODE): data portal update. Nucleic Acids Res, 46, D794–D801.

46. Barrett, T., Wilhite, S.E., Ledoux, P., Evangelista, C., Kim, I.F., Tomashevsky, M., Marshall, K.A., Phillippy, K.H., Sherman, P.M., Holko, M. et al. (2013) NCBI GEO: archive for functional genomics data sets--update. Nucleic Acids Res, 41, D991–995.

47. Narendra, V., Bulajic, M., Dekker, J., Mazzoni, E.O. and Reinberg, D. (2016) CTCF-mediated topological boundaries during development foster appropriate gene regulation. Genes Dev, 30, 2657–2662.

48. Nagano, T., Lubling, Y., Stevens, T.J., Schoenfelder, S., Yaffe, E., Dean, W., Laue, E.D., Tanay, A. and Fraser, P. (2013) Single-cell Hi-C reveals cell-to-cell variability in chromosome structure. Nature, 502, 59–64.

49. Yan, K.K., Yardimci, G.G., Yan, C., Noble, W.S. and Gerstein, M. (2017) HiC-spector: a matrix library for spectral and reproducibility analysis of Hi-C contact maps. Bioinformatics, 33, 2199–2201.

50. Serra, F., Bau, D., Goodstadt, M., Castillo, D., Filion, G.J. and Marti-Renom, M.A. (2017) Automatic analysis and 3D-modelling of Hi-C data using TADbit reveals structural features of the fly chromatin colors. PLoS Comput Biol, 13, e1005665.

